# Rejuvenation strategies share gene expression programs of reduced inflammation and downstream restored fatty acid metabolism

**DOI:** 10.1101/2022.09.26.509471

**Authors:** Tomer Landsberger, Ido Amit, Uri Alon

## Abstract

Understanding the mechanism of rejuvenation is central to aging research. No study has compared the effects of the four major rejuvenation strategies: senolytics, caloric restriction, *in vivo* partial cellular reprogramming and young/old blood factor exchange, which operate via different modalities. We use mice transcriptional data to compare them to each other and to normal aging. We find a shared gene expression program common to all rejuvenation strategies, in which inflammation declines and metabolism, especially of fatty acids, increases. An inverse signature occurs in normal aging. To test whether inflammation is upstream of the metabolic signature, we studied chronic inflammation in three different organs in young mice. Chronic inflammation was associated with a similar decline in metabolism, suggesting that inflammation is upstream of the metabolic signature. We find that inflammation may also underlie human transcriptional age calculator. We conclude that a core mechanism of rejuvenation acts through reduction of inflammation with downstream effects that enhance metabolism, attenuating the most robust age-related changes. This supports a notion of directly targeting genes associated with these pathways to mitigate age-related deterioration.

## Introduction

Aging is characterized by an accumulation of damage in multiple levels of biological organization that cause progressive loss of function and increased vulnerability to illness, thereby limiting organismal lifespan. Contrary to common perception, aging is not an inexorable entropic process. Multiple interventions operating via different modalities have been shown to delay and, in some cases, reverse age-related changes, and increase the lifespan of model organisms. *Mahmoudi et al*. 2019 ^1^ delineated four main strategies for organism rejuvenation and life-extension in mammals: 1. Exchange of blood-borne factors between young and old animals, e.g. by means of heterochronic parabiosis, a surgical procedure whereby two animals of different ages are joined together such that they share a common circulatory system. As a result of this exchange, young parabiont experience anti-aging effect whereas old parabiont experience pro-aging effects; 2. Metabolic manipulation, including caloric restriction, the oldest known and most widely applicable life-extending intervention; 3. Ablation of *senescence cells*, which are cells that underwent a stress-induced cell-cycle exit that accumulate with aging and secret pro-inflammatory agents; 4. *in vivo* partial cellular reprogramming, *in vivo* activation of Yamanaka factors in a transient way such that to achieve cellular rejuvenation while avoiding induction to pluripotent state and subsequent teratoma formation.

Despite extensive research into these interventions, and although translation efforts are already underway ^2^, key questions remain unanswered. A central question is what, if any, are the shared effects and mechanisms of action for these disparate interventions. Recently, extensive transcriptomic data pertaining to the four major rejuvenation strategies (RS) became available ^3–6^, calling for a comprehensive comparative analysis. Directly comparing the gene expression signatures of RS to one another and to that of normal aging may illuminate core aspects of rejuvenation and uncover new targets for therapeutics.

Here, we leverage bulk and single-cell RNA-seq (scRNA-seq) published datasets corresponding to the four RS, aging, and inflammatory disease models in mice, to investigate the shared effects and potential modes of action of RS. We identify a number of genes that are coordinately affected by RS in a pan-organ and cell-type manner. We find a robust RS pathway signature that coincides with the aging pathway signature, but in reverse. Inflammation and inflammation-related processes, as well as IGF transport and uptake, wound healing and cell-to-cell adhesion, are downregulated in all RS and upregulated in aging. Metabolic processes, and particularly lipid and fatty and carboxylic acid metabolism, are upregulated in all RS and downregulated in aging. By analyzing data from chronic inflammation in young mice, we show that metabolic changes we observed lay downstream of inflammation. Transcriptomic age calculator trained on human data assigns lower scores to samples from mice subjected to RS and higher scores to samples from chronic inflammation models, compared to their respective controls, suggesting inflammation to be a strong determinant of transcriptional age in mammals. Altogether, this study highlights the role of inflammation-related processes in rejuvenation and in normal aging.

## Results

### 1. Rejuvenation strategies share gene expression and pathway signatures with each other and with normal aging

We sought to compare the transcriptomic changes associated with the four major RS, to each other and to the transcriptomic changes associated with normal aging. To that end, we analyzed bulk and scRNA-seq data from four independent studies that tested these interventions in mice. 1. Ablation of senescent cells, achieved using the senolytic agent ABT-737 (ABT, *Ovadya et al*. ^3^). 2.Long-term caloric restriction (CR, *Rasa et al*. ^4^), decreased calorie intake without inducing malnutrition starting from 4 months of age. 3. *In vivo* partial reprogramming (IVPR, *Browder et al*. ^5^), achieved using transgene mice carrying a single copy of an OSKM polycistronic cassette which enable systemic expression of OSKM by administering doxycycline. 4. Heterochronic parabiosis (HP, *Ma et al*. ^6^), studied in multiple organs (of which we used liver-derived data) in single-cell resolution. Conveniently, each study also included young control to facilitate a direct comparison of the RS to aging. HP study included in addition to the heterochronic (Het) pair, old and young isochronic (Iso) pairs as controls.

Firstly, we re-derived ABT and IVRP-related differential expressed genes (DEGs) by comparing treated animals to their respective aged controls. Aged controls were also compared to young controls to generate intra-study aging reference signature. We used DESeq2 ^7^ standard workflow for these analyses, and considered genes with a |fold-change| > 1.25 and FDR < 0.05 to be differentially expressed (Supplementary table 1). For CR and HP, we used the DEGs data provided as supplementary for the original publication.

Next, we correlated the gene expression fold-changes (as calculated by DESeq2 or provided, hereinafter *signatures*) associated with RS in different organs with one another, and with the signatures associated with aging. For each pair, we used only the shared DEGs for the correlation, excluding pairs were shared DEGs set < 10. We performed this analysis separately for bulk and single-cell (HP) data.

We found that the RS signatures are mutually (Spearman) correlated in pan-organ and pan-intervention manner, and anti-correlated with the aging signatures, which are mutually correlated in pan-organ and pan-study manners (Fig. 1 A). For HP, analyzed on single-cell level, we find that Het-O/Iso-O (HP) signatures are correlated in a pan-cell-type manner, and anti-correlated with both Iso-O/Iso-Y (aging) and with Het-Y/Iso-Y (known to accelerate aging, hereinafter HP ACC) signatures. Consistent with that, aging and HP ACC signatures are mutually correlated in pan-cell-type manner (Fig. 1 A). We therefore conclude that RS partially reverse the aging signature in a similar way, and that HP ACC partially recapitulates the aging signature.

**Figure 1.**
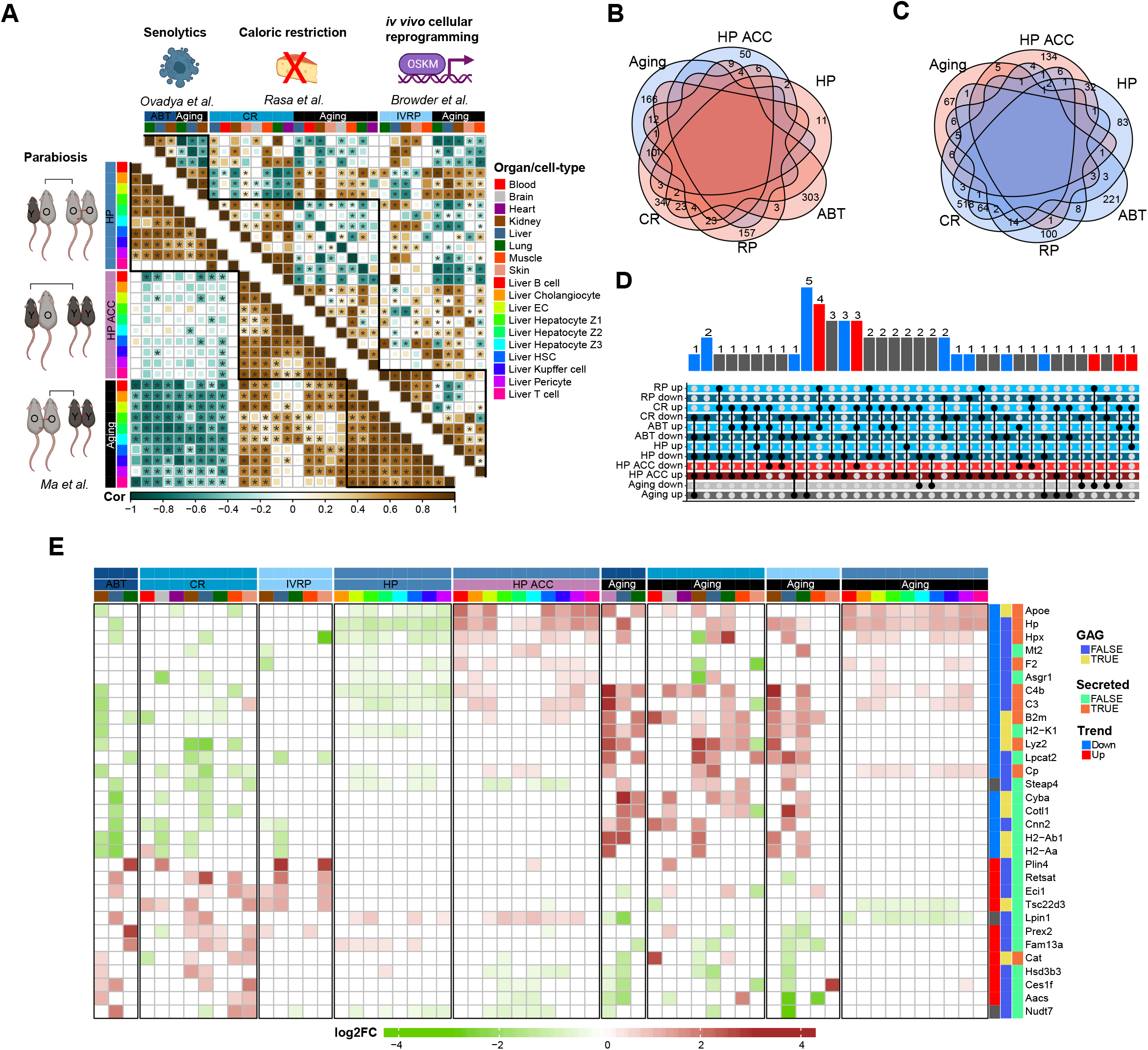
Rejuvenation strategies share a gene expression signature which is the inverse of aging: **A**. Spearman correlation of gene expression signatures of RS and same-study aging from different organ (top) and cell types (bottom). Only pair-wise correlations where shared DEG set > 10 are colored. Correlation coefficient encoded by both color and size. Asterisks depict p-value < 0.05. Color annotated for group and organ/cell type. **B**. Venn diagram of upregulated DEGs (fold-change > 1.25, p-value < 0.05) from ABT (>0 organs), CR (>1 organs), IVRP (>1 organs), HP (>2 cell types); and downregulated DEGs from HP ACC (>2 cell types) and aging (GAG). **C**. Same as B for downregulated RS DEGs and upregulated HP ACC DEGs and aging GAGs. **D**. Upset plot of 3^rd^ degree or greater intersects of DEG sets. Intersects consisting of downregulated RS and upregulated aging/HP ACC, or *vice versa*, are highlighted in blue/red respectively and numbered. **E**. heatmap depicting 3^rd^ degree or greater intersects of DEGs. Color coded for log2 expression fold-change (only DEGs, |fold-change| > 1.25, p-value < 0.05). Color annotated for study, group, and organ/cell type (up); and for trend (up/down in RS), GAG appearance, and if coding for secreted factor (right).

In order to identify genes that are robustly affected by RS and inversely by aging and HP ACC, we intersected the DEG sets derived from each of them. The aging-associated gene were imported from a study ^8^ that methodically analyzed the *Tabula Muris Senis* (TMS) ^9^, a comprehensive age-resolved murine bulk and single-cell transcriptomic dataset, to identify global aging genes (GAGs). GAGs consist of 93 upregulated and 190 downregulated genes. We intersected RS-associated upregulated DEGs with aging- and HP ACC-associated downregulated genes, and *vice versa* (Fig. 1 B, C). DEGs appear in up to 5 sets (Fig. 1 C). We examine the intersect distribution of DEGs that appear in more than 3 sets and find that, as expected, they tend to be either upregulated in RS and downregulated in aging and HP ACC, or *vice versa* (Fig. 1 D, highlighted in blue/red).

The DEGs that present this pattern (+3 DEGs that are close) are depicted in a heatmap (Fig. 1 E). The RS-downregulated genes consist mainly of immune-related genes. This demonstrates that RS reduce the low-grade inflammatory phenotype pervasive in mammalian aging (i.e., *inflammaging*). This set includes genes encoding for components of MHC class-I (*B2m, H2-K1*) and Class-II (*H2-Ab1, H2-Aa*) complexes, complement system (*C3, C4b*), and lysozyme (*Lyz2*). We also find *Apoe*, the gene whose polymorphisms are most associated with human longevity ^10,11^. Many of these genes encode for secreted factors, underscoring their potential systemic effect.

The RS-upregulated genes form a less coherent group in terms of function. Interestingly, *Cat*, which encodes for Catalase, is upregulated in both ABT and CR and downregulated in GAG. Catalase is a key antioxidant enzyme that defends against oxidative stress. It has been associated with age-related disease ^12^, and its mitochondrial ^13^ and cardiac ^14^ - targeted overexpression has a life-extending effect in mice. *Lpin1, Tsc22d3* and *Plin4* appear in the GenDR database ^15^, which curates genes whose expression is associated with dietary restriction. None of the other blue genes have previously been associated with RS or aging as per Human Aging Genomic Resources ^16^.

In order to characterize the biological processes that are commonly affected by the different RS, we applied gene set over-representation analysis by MetaScape ^17^ to multiple DEG sets derived from RS and aging in different organs and cell types. First, we co-analyzed downregulated RS DEGs with upregulated aging and HP ACC DEGs. We scored each pathway according to the number of tested DEG sets where it is enriched. Each whole organ-related hit contributed 1 point and each cell type-related hit from HP study contributed 0.5 point to total rank score. We found that the RS/HP ACC and aging rank scores are correlated, and particularly that the highest scoring pathways for RS/HP ACC are also the highest scoring pathways for aging (Fig. 2 A). This means that the shared RS downregulated pathways signature coincides with the aging upregulated pathways signature.

**Figure 2.**
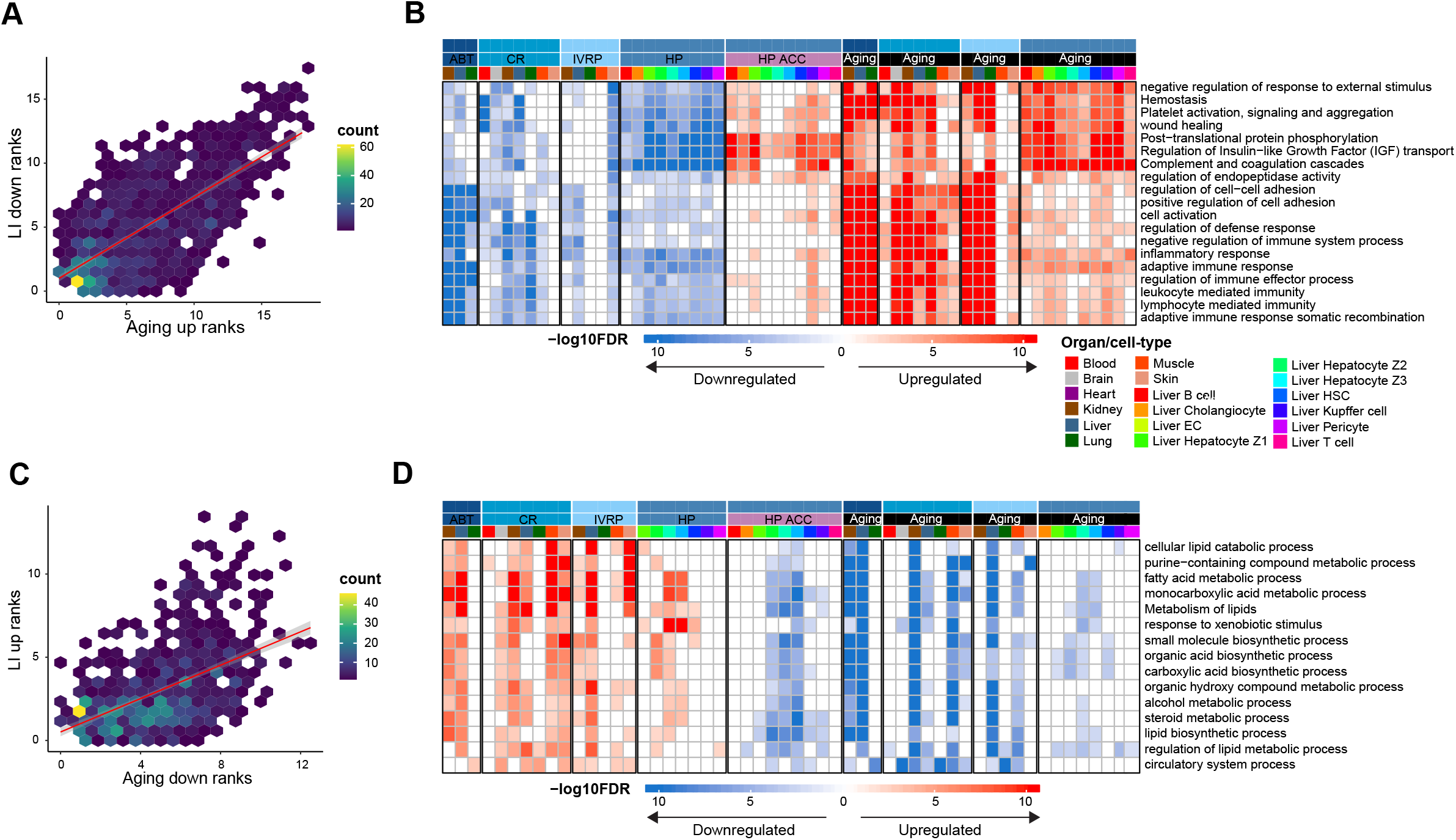
Rejuvenation strategies share pathway signature which is the inverse of aging: **A**. aging upregulated pathways rank scores (x axis) vs RS downregulated/HP ACC upregulated pathways rank scores (y axis), calculated by the number of DEG sets where a pathway is enriched (1 point for whole organ and 0.5 point for cell type). **B**. heatmap depicting top scoring pathways for RS down/HP ACC up, alongside aging. Color coded for -log10 FDR, with blue downregulated and red upregulated, showing only FDR < 0.01. Color annotated for study, group and organ/cell type. **C**. same as A but for opposite directions. **D**. same as B. for opposite directions.

Some of the most robust shared pathways were inflammation-related, including complement system and adaptive immune responses (Fig. 2 B, Supplementary table 2), consistent with the gene-level analysis. Inflammation has been shown to constitute a gene expression hallmark of aging in different organs ^18^ and mammalian species ^19^. With respect to RS, *Mahmoudi et al*.^1^ concluded that inflammation is mitigated in CR and HP, but that only indirect evidence exists to support its mitigation in senolytics and IVRP. Here we observe a ubiquitous mitigation of inflammation by RS.

Other interesting pathways are wound healing, cell-to-cell adhesion, and regulation of Insulin−like Growth Factor (IGF). The latter pathway is tightly linked to aging, and its inhibition in adult or late life confers life-extension in female mice ^20^. Lower scoring pathways also included neutrophil degranulation, ERK1/2/MAPK cascade, phagocytosis, and ECM organization (Supplementary table 2).

We applied the same analysis to DEG sets that are upregulated in RS and downregulated in aging and HP ACC. Here too a correlation exists between RS/HP ACC and aging related pathways (Fig. 2 C). The pathways most strongly associated with these sets are metabolism-related, especially lipid, and fatty and carboxylic acid metabolism (Fig. 2 D, Supplementary table 2). This is line with diverse evidence suggesting that lipid metabolism is an important regulator of aging in nematodes, fruit flies, mice, and rats ^21^. Moreover, several other studies conducting transcriptomic analysis have reported changes in these pathways in the context of aging ^22^ (reduced), cellular senescence^23,24^ (enhanced), HP ^25^ (enhanced), and especially in the context of CR ^18,26–28^ (enhanced), where it is highly foreseeable. Recently, the senolytic agents dasatinib and quercetin were shown to have similar effects on both inflammation and metabolism in adipose tissue, using targeted assays ^29^.

Our analysis shows that a reduction of inflammation and an increase fatty acid metabolism occur in all RS, implying a mechanistic link may exist between these processes. To test this in a non-aging context, we sought to study the transcriptomic signature of inflammation in young animals.

### 2. Chronic inflammation-eliciting disease models in young mice show that metabolic changes lay downstream of inflammation

We analyzed both bulk and scRNA-seq data derived from murine models of three diseases that involve chronic inflammation (hereinafter referred to as CI models) – idiopathic pulmonary fibrosis (IPF. bulk: *Matsuda* et al. ^30^, single-cell: *Strunz* et al. ^31^), non-alcoholic steatohepatitis (NASH, bulk: *Xiong* et al. ^32^, single-cell: *Xiong* et al. ^33^), and obstructive nephropathy (ON, bulk: *Arvaniti* et al. ^34^, single-cell: Conway et al. ^35^). To establish an aging reference, we used organ- and technology-matched TMS data, with the exception of single-cell lung, where we used data from *Angelidis* et al. ^36^ lung atlas (Methods).

For bulk datasets, we re-derived DEGs using DESeq2. For single-cell datasets, we re-derived DEGs and in some cases also the cell clustering and annotation, using Seurat ^37^ package suite and MAST algorithm ^38^ for detection of DEGs, implemented thereof (Supplementary table 1, Supplementary Fig. 1, methods).

Following the same protocol as for RS and aging, we pairwise (Spearman) correlated the gene expression signatures of CI to each other and to aging, separately for bulk and single-cell data, revealing pan-organ and pan-cell-type correlations respectively (Fig. 3 A). This shows that the three CI models analyzed partially recapitulate the gene expression signature of aging.

**Figure 3.**
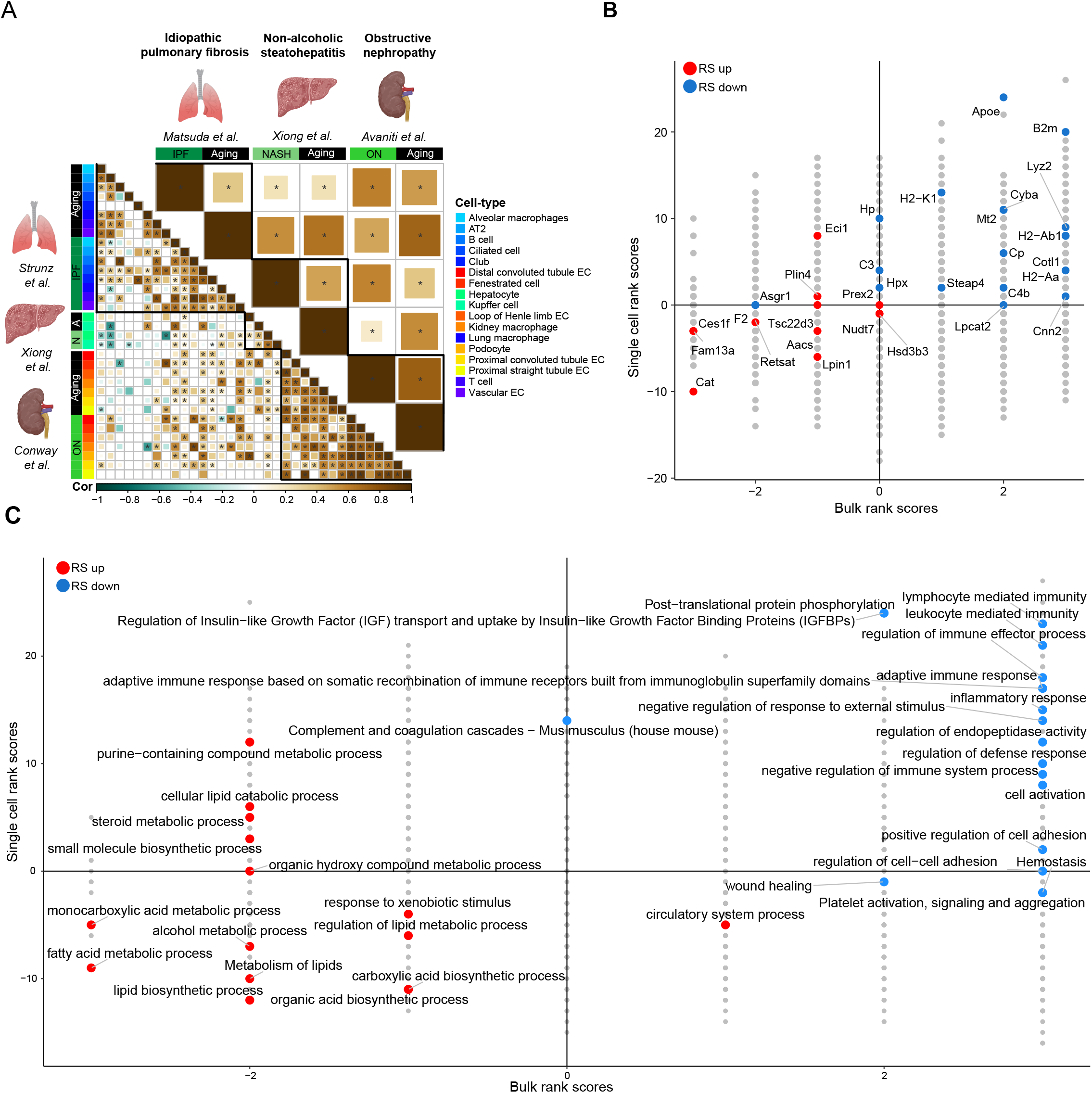
Age-related metabolic changes reversed by rejuvenation strategies are downstream of inflammation: **A**. Spearman correlation of gene expression signatures for CI models and organ-matched aging, in bulk (top) and single-cell (bottom) levels. Only pair-wise correlations where shared DEG set > 10 are colored. Correlation coefficient encoded by both color and size. Asterisks depict p-value < 0.05. Color annotated for group and cell type. **B**. 2d map of CI-related DEG rank scores in bulk (x axis) and single-cell (y axis) levels. Score calculated by number of organs (bulk) or cell types (single-cell) where gene is a DEG (|fold-change| > 1.25, p-value < 0.05). Upregulation scores 1 point and downregulation scores −1 points. RS-related DEGs from Fig. 1 E are highlighted in color, in accordance to their trend in RS. **C**. 2d map of CI-related pathway enrichment rank scores in bulk (x axis) and single-cell (y axis) levels. Upregulation scores 1 point and downregulation scores −1 points. RS-related pathways from Fig. 2 B and D are highlighted in color, in accordance to their trend in RS.

Next, we scored each gene according to the number of times it is significantly upregulated (FC > 1.25, FDR < 0.05) in one organ (x-axis) or cell type (y-axis) in the CI models data, minus the number of times it is downregulated (Fig. 3. B). Highlighting the RS-shared DEGs presented in Figure 1E on that map of CI-scored DEGs, we find that many of them are differentially expressed in several tissue and cell types, in the opposite direction to RS. *B2m* was the most robust upregulated gene, and *Cat* the most robust downregulated gene. The downregulation of *Cat, Fam13a* and *Ces1f* in all three CI models suggest that their upregulation in RS is downstream of inflammation.

We used MetaScape ^17^ to analyze the DEG sets associated with CI models in different organs and cell types. We scored each pathway in the same way as for genes, counting enrichment (FDR < 0.01) in a given set of DEGs instead of differential expression. Highlighting the RS-associated pathways from Figure 2 B and D on the CI-scored enriched pathways map, we find that inflammation-related pathways are predictably upregulated (Fig. 3 C, Supplementary table 2). Wound healing, and cell-to-cell adhesion are upregulated only in bulk level. Interestingly, metabolism-related pathways are robustly downregulated. Fatty acid metabolism is the most consistently downregulated pathway across CI models. These results indicate that the observed metabolic changes occur downstream of inflammation.

### 3. Transcriptional age is affected by inflammation

Several studies have reported the accelerating or deaccelerating effect of different interventions on so-called *biological age*, as measured by multiparametric *gerometers* (AKA aging clocks), most commonly DNA methylation markers ^39^. We hypothesize that inflammation-related signals, comprising the main shared component of both RS and aging gene expression signatures, are strong determinants of biological age as measured by these methods. To test this, we used RNAAgeCalc ^40^, calibrated on human data (whole organ bulk RNA-seq) curated in GTEx ^41^ data repository. Mouse gene names were converted to their human homologs (methods) to enable this cross-species application, and the age readout was scaled.

We find that RNAAgeCalc assigns higher RNA age to samples from old (24/27 months) compared to young (3 months) mice from TMS (Fig. 4 A). Furthermore, in many cases, RS reduce RNA age compared to aged controls (Fig. 4 B). IVPR reduces RNA age to a significant level only in skin (Fig 4. B), consistent with original study reporting significant reduction in DNA methylation exclusively in that tissue ^5^. Most importantly, in agreement with our hypothesis, RNAAgeCalc assigns significantly higher RNA age to samples from inflamed organs compared to their respective controls (Fig. 4 C). Altogether, this demonstrates that inflammation-related genes constitute strong pan-species (mammalian) aging biomarkers that potentially underlie multiparametric gerometers.

**Figure 4.**
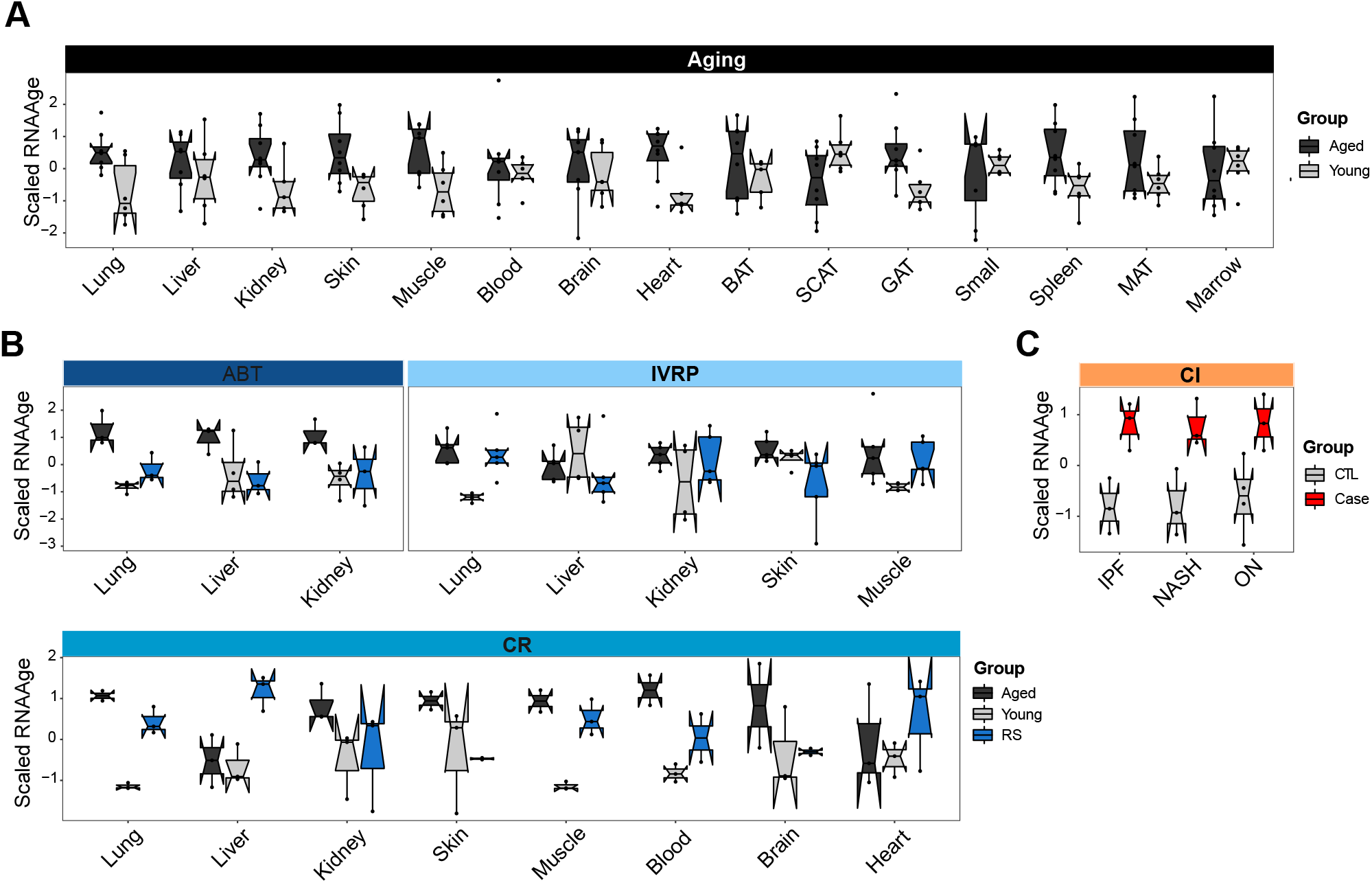
Transcriptional age is affected by inflammation: **A**. Scaled RNA age (y axis) as calculated by RNAAgeCalc for TMS-derived bulk gene expression profiles of old (24/27 months old) and young (3 months old) mice in different tissues (x axis). **B**. Scaled RNA age (y axis) for ABT, CR and IVRP bulk gene expression profiles in different organs (x axis). **C**. Scaled RNA age (y axis) for IPF, NASH and ON bulk gene expression profiles. Points represent samples in all panels. Boxplot centers represent median values. Color coded for group.

## Discussion

Recent years have seen a surge in attention given to RS, with both academic research ^1^ and biotech spin-off companies ^2^ taking notice. Despite the increased focus, our understanding of the mechanisms and effects of RS remains limited, calling for further basic scientific exploration. In this study, we reveal a shared feature among the four major RS: a reduction in inflammation. This decrease in inflammation also leads to the restoration of fatty acid metabolism, which is diminished during normal aging. While the role of chronic inflammation in aging is well-known ^18,42,43^, its role in RS has been less widely acknowledged. Our findings suggest that inflammation and related metabolic changes should be considered hallmarks of not only aging but also rejuvenation.

Our understanding of the aging process implies that changes in gene expression as a result of aging are a combination of passive, aging-promoting changes caused by a drift in the transcriptional landscape and active, aging-antagonizing changes brought on by stress response mechanisms aimed to counteract various damage types incurred to the organism and restore homeostasis. Although these two classes of changes can be difficult to separate, we generally expect the latter to be more coordinated and coherent than the former. Inflammation is a clear example of a programmed response meant to counteract damage to the organism, but it can become deleterious if it persists over time. The metabolic changes we observed in our study may be a less known result of this harmful overshooting of inflammation, i.e., inflammaging. Alleviating inflammaging and restoring proper metabolism could be one way in which RS exsert their anti-aging effects.

RS may alleviate inflammaging directly, or indirectly by removing the damage signals that cause it. Senolysis eliminates senescent cells thereby reducing SASP (senescence-associated secretory phenotype) that involves many inflammatory signals ^44^; Rasa *et al*. suggested CR curbs inflammation by reducing chromatin accessibility of an inflammation-associated genetic network^4^. IVPR has been shown to have a rejuvenative effect both *ex vivo*^45^ and *in vivo*^46^, which could involve deactivating stress signals from damaged cells. Lastly, HP may reduce systemic inflammation by diluting pro-inflammatory mediators in old blood or introducing youthful anti-inflammatory factors. This has been shown by Jeon *et al*. who reported that senolytic treatment of old donors reduced pro-aging effects in young recipients after blood exchange ^47^.

The opposite effects CI models and RS have on transcriptional age are aligned with reported effects on DNA methylation (DNAm) age ^39^. Studies have shown that inflammatory conditions such as Covid-19 infection ^48^ can accelerate DNAm age, while lifestyle changes ^49^, plasma fraction treatment^50^, cellular reprogramming ^51^, HP ^25^, or drug cocktail ^52^, can reverse it. Evidence also suggests that even a transient inflammatory stimulus can accelerate DNAm age and impair self-renewal in hematopoietic stem cells ^53^. These findings, in conjunction with our results, raise the possibility that inflammatory signals contribute significantly to readings of widely used multiparametric aging gerometers. That may be due to strong signals of inflammatory processes, compared to potentially more subtle and diffused, yet also more stable, age-related signals. Further research is needed to understand the role of inflammation and downstream signaling in these readings.

In conclusion, our analysis points to inflammation and reduced fatty acid metabolism as robust hallmarks of aging that are reversed by the four major RS. A possible interpretation of our findings is that these two hallmarks are important mediator of the health benefits of said RS. This would argue in favor of directly targeting inflammation and fatty acid metabolism pathways to mitigate age-related health decline.

## Abbreviations

TMS: Tabula Muris Senis
scRNA-seq: Single-cell RNA sequencing
DEG: Differentially expressed gene
RS: Rejuvenation strategies
ABT: ABT-737 mediated senolytic treatment
CR: Caloric restriction
IVPR: *in vivo* partial reprogramming
HP: Heterochronic parabiosis
HP ACC: Heterochronic parabiosis-related accelerated aging
CI: Chronic inflammation.
IPF: Idiopathic pulmonary fibrosis NASH – Non-alcoholic steatohepatitis
ON: Obstructive nephropathy
EC: Endothelial cell
DNAm: DNA methylation

## Methods

### Choice of datasets for analysis

RS datasets were chosen base on availability, breath, and quality of publication. Aging datasets used as reference in the analysis of CI models were taken from TMS based on the comprehensiveness of this database. Droplet data was used for optimal match with CI dataset. For the analysis of single-cell aging lungs, *Angelidis* et al. aging lung cell atlas ^36^ was used instead of TMS owing to greater cell-type overlap with *Strunz* IPF dataset. In IVRP, spleen samples were excluded for poor signal.

### Bulk differential gene expression analysis

the standard workflow of DESeq2 ^7^, which is based on the negative binomial distribution, was used for all bulk datasets. DGEs are selected to have |fold-change| > 1.25 and Benjamini-Hochberg adjusted p-value < 0.05. Tests were controlled for sex as a covariant. For TMS data, 24- and 27-month-old mice were compared to 3-month-old mice. For CR, DEG lists provided in the original publication were used.

### Sc clustering and cell-population annotation

to facilitate direct comparison of CI models to aging from independent studies, some of the annotations that appeared in the original publications needed to be re-established so that cell-types can be accurately matched (Supplementary Fig. 1). Cell type annotations were adopted from original studies when applicable (lung: *Strunz/Angelidis*, kidney: TMS). Where published annotation was deemed too coarse-grained (lung: TMS, liver: *Xiong*/TMS, kidney: *Conway*), Seurat package (version 4.0) ^37^ standard workflow was used for clustering. Briefly, cells with > 15% mitochondrial content were removed, UMI matrix was log-normalize, highly variable genes were detected (following exclusion of mitochondrial, ribosomal and cell-cycle genes), and PCA was applied, followed by KNN graph construction on 50 leading components. Louvain algorithm was used for community detection using resolution parameter 1.5. Clusters were manually annotated based on marker genes and cross-reference with the pre-annotated dataset. Note that not all mice used for clustering are used for downstream analysis.

### Sc ambient RNA removal

ambient RNA is RNA present in the cell suspension that can be aberrantly counted along with a cell’s native mRNA. In most datasets, ubiquitous expression of highly-expressed cell-type specific genes (e.g., surfactant genes in lung tissue), suggested such cross-contamination. To overcome this, decontX() function from celda (V1.6.1) ^54^ was used with default parameters to remove contamination in individual cells. decontX is a Bayesian method that models the empirical expression of a cell is a mixture of counts from two multinomial distributions: (1) a distribution of native transcript counts from the cell’s actual population and (2) a distribution of contaminating transcript counts from all other cell populations captured in the assay.

#### Sc differential gene expression analysis

For HP data, DEGs were adopted from original study. For CI models data, and for TMS-derived aging reference, DEGs were re-derived, following clustering and annotations (as per Sc clustering and cell-population annotation), using two-sided MAST ^38^ test implemented in Seurat ^37^ V4. MAST analysis was applied to decontaminated log-transformed normalized counts for each group and each cell-type individually. Genes were pre-filtered to those expressed in > 0.01 of cells and with > 5 cells with > 1 UMIs. DGEs are selected to have |fold-change| > 1.25 and Benjamini-Hochberg adjusted p-value < 0.05. Tests were controlled for sex as a covariant.

In order to avoid ambient RNA, in addition to using DecontX corrected data, we also excluded DEGs if they met both these criteria: 1. among the top 5 markers in another cell type *and* 2. among the top 50 most highly expressed genes in the dataset.

We note that for *Strunz* et al. lung fibrosis study, who performed timecourse measurements, we used cells from 7 days post-bleomycin treatment onwards.

For liver and kidney from TMS, tests were applied for aged vs. young, where “aged” consists of cells pooled from 18-, 21-, 24- and 30-month-old mice, and “young” consisted of 3-month-old mice.

#### Gene set over-representation analysis

MetaScape ^17^ web interface for multiple gene lists was used to analyze DEGs with parameters: Min Overlap = 2, P Value Cutoff = 0.01, and Min Enrichment = 1.5.

#### Correlation analysis

we performed pair-wise Spearman correlations of the fold-changes of shared DEGs for each pair. When DEG set was < 10, the pair was excluded. Only correlations where p-value < 0.05 are presented in the heatmaps.

### Application of RNAAgeCalc gerometer

RNA age was calculated using RNAAgeCalc ^40^ trained on human data derived from GTEx V6 to construct a cross-tissue and tissue-specific transcriptional age calculator. The algorithm was implemented in R following the guidelines described in https://www.bioconductor.org/packages/release/bioc/vignettes/RNAAgeCalc/inst/doc/RNAAge-vignette.html, with tissue specific predictor used when applicable and pan-tissue predictor used for liver and kidney and muscle. The counts matrix was supplied. When counts were UMIs (ABT), gene lengths were set to 10K for all genes. Mouse gene names were converted to human homologs using convert_mouse_to_human_symbols() function from nichenetr ^55^ library in R. The readout, given in human years, was scaled for presentation.

### Visualization

Plots were generated in R using ggplot2 ^56^, ComplexHeatmap ^57^, UpSetR ^58^, and corrplot ^59^ R libraries.

## Acknowledgments

We are grateful to Lorenz Adlung and Amir Giladi for providing helpful comments.

## Authors’ contributions

T.L. performed data analysis. T.L. and U.A. design the research. T.L. and U.A. wrote the manuscript. U.A. and I.A. supervised the study.

## Supplementary material

Supplementary material for this study can be found online.

