## Supplementary Figure 1 for "Rejuvenation strategies share gene expression programs of reduced inflammation and downstream restored fatty acid metabolism"

### Supplementary Figure 1 - ScRNA-seq clustering and cell type annotation for CI models and aging

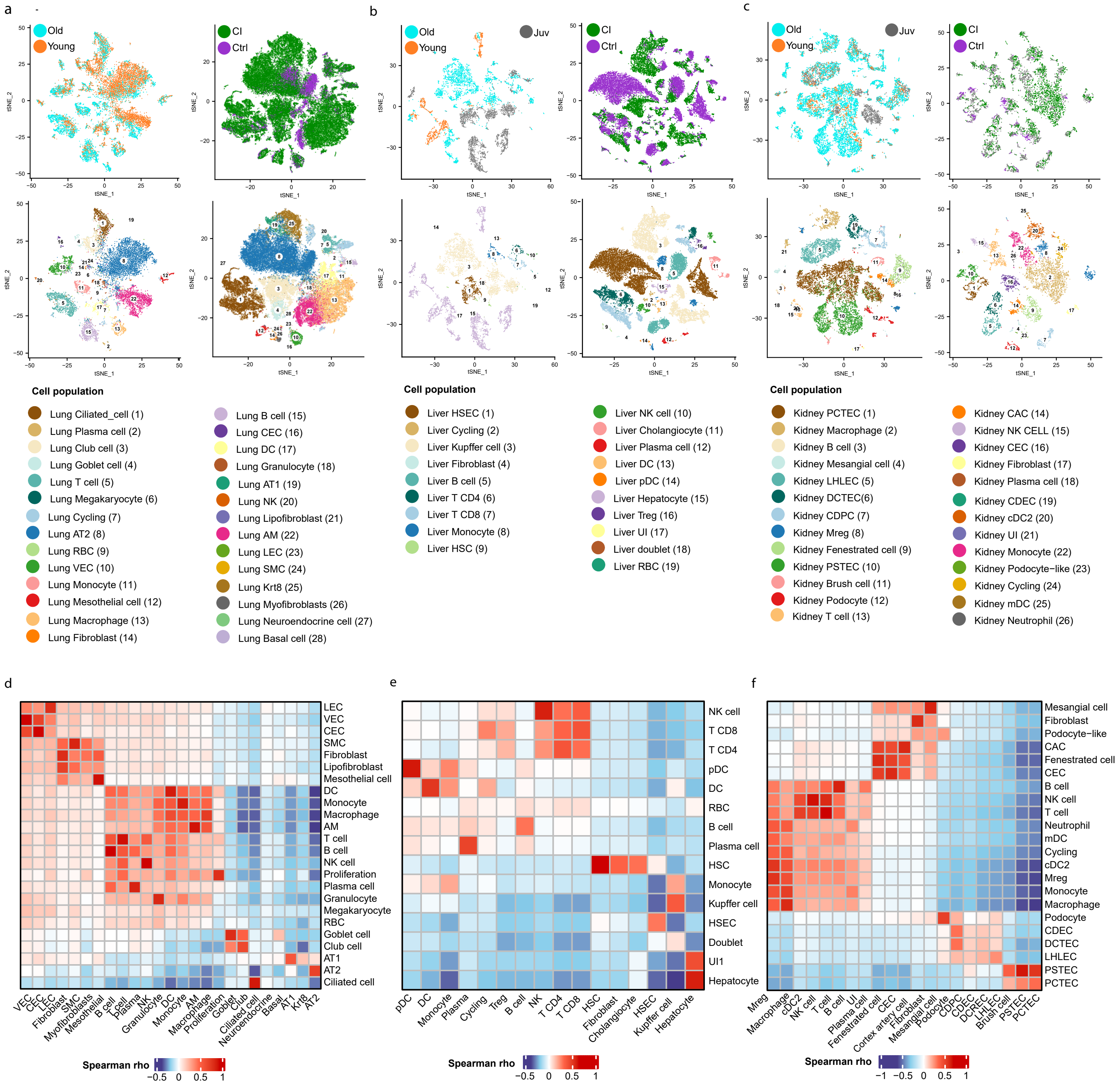


**Supplementary Figure 1 - ScRNA-seq clustering and cell type annotation for CI models and aging.**
a-c, t-distributed stochastic neighbor embedding (tSNE) maps of single cells from aging (left) and CI (right) studies analyzed, color coded for experimental group (top) and for cell population (bottom), in lung (a), liver (b) and kidney (c). d-f, Inter-study Spearman correlation per cell population of top 3000 most variable genes in lung (d), liver (e) and kidney (f).
